# Disrupting the leukemia niche in the central nervous system attenuates leukemia chemoresistance

**DOI:** 10.1101/750547

**Authors:** Leslie M. Jonart, Maryam Ebadi, Patrick Basile, Kimberly Johnson, Jessica Makori, Peter M. Gordon

**Affiliations:** Division of Pediatric Hematology and Oncology, Department of Pediatrics, University of Minnesota, Minneapolis, MN, USA; Masonic Cancer Center, University of Minnesota, Minneapolis, MN, USA

## Abstract

Protection from acute lymphoblastic leukemia (ALL) relapse in the central nervous system (CNS) is crucial to survival and quality of life for ALL patients. Current CNS-directed therapies cause significant toxicities and are only partially effective. Moreover, the impact of the CNS microenvironment on leukemia biology is poorly understood. Herein, we showed that leukemia cells associated with the meninges of xenotransplanted mice, or co-cultured with meningeal cells, exhibit enhanced chemoresistance due to effects on both apoptosis balance and quiescence. From a mechanistic standpoint, we identified that leukemia chemoresistance is primarily mediated by direct leukemia-meningeal cell interactions and overcome by detaching the leukemia cells from the meninges. Next, we used a co-culture adhesion assay to identify drugs that disrupted leukemia-meningeal adhesion. In addition to identifying several drugs that inhibit canonical cell adhesion targets we found that Me6TREN, a novel hematopoietic stem cell (HSC) mobilizing compound, also disrupts leukemia-meningeal adhesion *in vitro* and *in vivo*. Finally, Me6TREN enhanced the efficacy of cytarabine in treating CNS leukemia in xenotransplanted mice. This work demonstrates that the meninges exert a critical influence on leukemia chemoresistance, elucidates mechanisms of CNS relapse beyond the well-described role of the blood-brain barrier, and identifies novel therapeutic approaches for overcoming chemoresistance.

## Introduction

Central nervous system (CNS) relapse is a common cause of treatment failure among patients with acute lymphoblastic leukemia (ALL)^1–3^. Relapses occur despite CNS-directed therapies which include high-dose systemic chemotherapy, intrathecal chemotherapy, and cranial irradiation in some high-risk patients. These current CNS-directed therapies are also associated with significant acute and long-term toxicities^4–10^. Accordingly, novel CNS-directed leukemia therapies are needed to improve long-term outcomes in ALL while decreasing treatment-related morbidity.

Historically, the ability of leukemia cells and chemotherapy to access the restricted CNS environment has been posited as critical factors in the pathophysiology of CNS leukemia and relapse. However, several lines of evidence suggest this is an overly simplistic model. First, high rates (>50%) of CNS leukemia occur in patients in the absence of adequate CNS-directed therapies as well as in mice transplanted with human, primary B-cell precursor leukemia cells^11–14^. Moreover, clonal analyses of paired leukemia cells isolated from both the bone marrow and CNS of patients and xenotransplanted mice demonstrated that all, or most, B-cell ALL clones are capable of disseminating to the CNS^14,15^. Third, CNS leukemia relapses occur despite high dose systemic and intrathecal chemotherapy. These therapies either overcome or bypass the blood-brain barrier. Fourth, it was shown that high Mer kinase expressing, t(1;19) leukemia cells co-cultured with CNS-derived cells exhibit G0/G1 cell cycle arrest, suggestive of dormancy or quiescence, as well as methotrexate resistance^16^. Similarly, Akers *et al*. showed that co-culture of leukemia cells with astrocytes, choroid plexus epithelial cells, or meningeal cells enhanced leukemia chemoresistance^17^. Together these observations suggest that the pathophysiology of CNS leukemia extends beyond the role of the blood-brain barrier. We hypothesize that the ability of leukemia cells to persist in the unique CNS niche and escape the effects of chemotherapy and immune surveillance likely also play critical roles in CNS leukemia and relapse.

However, while extensive research has demonstrated a critical role of the bone marrow niche in leukemia biology, the impact of the CNS niche on leukemia biology is less well understood^18,19^. Herein, we demonstrate that the meninges exert a unique and critical influence on leukemia biology by enhancing leukemia resistance to the chemotherapy agents currently used in the therapy of CNS leukemia. We then leveraged this new understanding of the mechanisms of meningeal-mediated leukemia chemoresistance to identify a novel drug, Me6TREN (Tris[2-(dimethylamino)ethyl]amine), which overcomes leukemia chemoresistance by disrupting the interaction between leukemia and meningeal cells.

## Materials & Methods

### Cells and tissue culture

Leukemia cells were obtained from American Type Culture Collection (ATCC) or DSMZ and cultured in RPMI media (Sigma-Aldrich) supplemented with Fetal Bovine Serum (FBS; Seradigm) 10% and Penicillin-Streptomycin (Sigma-Aldrich). Leukemia cell lines included both B- (NALM-6, SEM) and T-cell (Jurkat, SEM, MOLT-13) immunophenotypes. The HCN-2 neuronal cell line was obtained from ATCC. Leukemia cells expressing green fluorescent protein (GFP) were generated as described^20^. Murine leukemia cells, generated by BCR/ABL p190 expression in hematopoietic cells from CD45.1 Arf^-/-^ mice^21–23^, were provided by Dr. Michael Farrar (University of Minnesota). Primary B-ALL cells for co-culture experiments were obtained from the University of Minnesota Hematological Malignancy Bank (IRB #: 0611M96846; pediatric patient at diagnosis). Primary B-ALL cells for *in vivo* experiments were obtained from the Public Repository of Xenografts (PRoXe^24^; Sample CBAB-62871-V1; pediatric patient at diagnosis with a t(4;11) translocation). Primary meningeal cells were obtained from ScienCell and cultured in meningeal media supplemented with FBS 2%, growth supplement, and Penicillin-Streptomycin. Meningeal cells were isolated from multiple different donor specimens and were typically used between passages 3-5.

### Leukemia co-culture

Leukemia cells were plated onto ~60-70% confluent primary meningeal cells and grown in a mixture of RPMI and meningeal cell media with FBS 10%. Non-adherent leukemia cells were removed by aspirating the media after overnight co-culture. Fresh media was then added and the cells were co-cultured for 24-48 hours in the presence or absence of drugs. Drugs, including cytarabine, methotrexate, and Me6TREN, were obtained from Sigma-Aldrich. Drug doses were based on the literature^17,25–28^ and drug titrations that identified concentrations of cytarabine and methotrexate that caused significant apoptosis to leukemia cells in suspension. After the specified co-culture period, media containing non-adherent leukemia cells was collected. Next, leukemia cells were removed from adhesion to meningeal cells with 0.05% Trypsin and manual pipetting. Leukemia cells were combined and assessed for apoptosis and cell cycle by flow cytometry as described. During flow cytometry, leukemia cells were distinguished from any contaminating meningeal cells by staining with leukemia-specific fluorescent antibodies (CD19, CD3, CD45; eBioscience) or the leukemia cells were fluorescently labeled with CellTrace dye (ThermoFisher) immediately prior to co-culture. For some experiments, the leukemia cells were isolated from meningeal cells after co-culture using immunomagnetic separation with either CD19 or CD3 antibodies (StemCell Technologies or Miltenyi Biotec).

### Apoptosis and cell cycle analyses

To assess apoptosis, leukemia cells were treated as described, stained with annexin-V antibody (eBioscience) and a fixable viability dye (ThermoFisher) or 7-AAD, and analyzed by flow cytometry using a BD FACSCanto. TMRE (Abcam) and Caspase 3/7 assays (ThermoFisher) were performed according to the manufacturer’s instructions. The Human Apoptosis Antibody Array (Abcam; 43 targets) was performed with protein lysates from leukemia cells grown in suspension or isolated from co-culture using immunomagnetic separation. Cell cycle and proliferation were assessed using a Click-iT Plus EdU flow cytometry kit (ThermoFisher). To differentiate G0 from G1 phases of the cell cycle, leukemia cells were stained with both Hoechst 33342 (Sigma-Aldrich) and pyronin Y (Sigma-Aldrich) and analyzed by flow cytometry^29^. To assess long-term quiescence *in vivo*, immediately prior to transplantation leukemia cells were labeled with the membrane dye DiR (1,1’-Dioctadecyl-3,3,3’,3’-Tetramethylindotricarbocyanine Iodide; ThermoFisher) for microscopy or DiD (1,1’-dioctadecyl-3,3,3’,3’-tetramethylindodicarbocyanine, 4-chlorobenzenesulfonate salt; ThermoFisher) for flow cytometry^28,30–33^. Ki-67 levels were measured with a fluorescent antibody (eBioscience) and flow cytometry.

### Murine experiments

NSG (NOD.Cg-*Prkdcscid, Il2rgtm1Wjl*/SzJ; Jackson Labs) mice were housed under aseptic conditions. Mouse care and experiments were in accordance with a protocol approved by the Institutional Animal Care and Use Committee at the University of Minnesota (IRB#1704-34717A). Mice 6-8 weeks old were injected intravenously via the tail vein with ~1-2×10^6^ human leukemia cells. Experiments were then performed 3-5 weeks after injection. In general, after euthanasia the mice were cardiac perfused with PBS, meninges removed using a dissecting microscope, and dissociated by gently washing through a 0.40 μm filter (Millipore). Cells were then stained with fluorescent antibodies against CD19 (NALM-6; eBioscience) or CD3 (Jurkat, eBioscience) and assessed for apoptosis or cell cycle as described. Alternatively, leukemia cells could be purified from meningeal cells using immunomagnetic separation and either CD19 or CD3 antibodies (Stem Cell Technologies or Miltenyi Biotec) and placed back into suspension. In drug treatment experiments, Me6TREN 10 mg/kg and cytarabine 50 mg/kg were given by subcutaneous and intraperitoneal injection, respectively. Cerebral spinal fluid (CSF) was obtained from mice as described^34^. Briefly, mice were euthanized and a scalp incision was made at the midline to expose the dura mater overlying the cisterna magna. Under a dissection microscope, a tapered, pulled glass capillary tube was then inserted through the dura and into the cisterna magna to obtain clear cerebral spinal fluid (CSF). Experiments with murine leukemia cells (BCR/ABL p190 expression in hematopoietic cells from Arf^-/-^ mice; CD45.1 background) used C57BL/6 mice. In these experiments, ~3000 leukemia cells/mouse were injected via the tail vein.

### Microscopy

Leukemia cells stably expressing GFP were labeled with the fluorescent membrane dye dIR (ThermoFisher) immediately prior to xenotransplantation^30,31^. At signs of systemic leukemia, mice were euthanized and cardiac perfused with PBS followed by paraformaldehyde 4% (pH 7.4). Skin, subcutaneous tissue, nasal bones and the mandible were then removed. The cranium was decalcified using a Pelco BioWave Pro Processor^35^ and then the skull, brain and intact meninges were sectioned with a vibratome at ~600 μm per section. Sections were re-processed in a Pelco BioWave Pro Processor to improve depth of imaging and reduce light scatter. Meningeal sections were imaged using a Nikon A1R FLIM Confocal Microscope using the 488 nM and 640 nM laser for GFP and DiR detection, respectively. Images were analyzed using ImageJ software.

### Statistical analysis

Results are shown as the mean plus or minus the SEM of the results of at least 3 experiments. The Student’s *t*-test or ANOVA were used for statistical comparisons between groups and were calculated using GraphPad Prism 7 software (GraphPad Software, La Jolla, CA). *P*-values < 0.05 were considered statistically significant.

## Results

### Leukemia cells reside in the meninges of the mouse CNS

In order to identify the anatomic site(s) within the CNS that leukemia cells reside, we transplanted multiple human ALL cell lines, including NALM-6, Jurkat, and SEM, into immune-compromised mice (NSG) via tail vein injection (Supplemental Figure 1A). Mice were not irradiated or conditioned with busulfan prior to transplantation to avoid perturbing leukemia niches. The mice were then euthanized and the CNS examined by histopathology and immunohistochemistry. We identified both the meninges and, to a lesser extent, the choroid plexus as the predominant CNS sites that harbor leukemia cells both before and after treatment with systemic cytarabine (Supplemental Figure 1B). In contrast, parenchymal involvement by leukemia was a rare, and often late, finding. It is possible that the altered immune system of NSG mice could influence CNS leukemia involvement or anatomic distribution. Accordingly, we also tested a pre-B ALL mouse leukemia model that utilizes BCR/ABL p190 expression in hematopoietic cells from *Arf^-/-^* mice transplanted into immunocompetent mice^21,22,36^. Similar to the xenotransplantation results, leukemia extensively involved the meninges in these mice (Supplemental Figure 1C).

### The meninges enhance leukemia chemoresistance

We then developed *ex vivo* co-culture approaches to focus more specifically on the effects of the meninges on leukemia chemosensitivity. We selected meningeal cells based on our IHC analyses of brains from transplanted mice (Supplemental Figure 1B-C) as well as histopathological examinations of brains from leukemia patients^11–14^. We found that leukemia cells adhered to primary human meningeal cells in a co-culture system (Figure 1A). Moreover, leukemia cells co-cultured with meningeal cells were significantly more resistant to cytarabine and methotrexate-induced apoptosis, as measured by annexin-V and viability staining, relative to the same cells grown in suspension or adherent to the HCN-2 neural precursor cell line (Figure 1B and Supplemental Table 1). In these experiments, chemotherapy had only very modest effects on meningeal cell viability (Supplemental Figure 2). Additionally, primary pre-B leukemia cell survival both in the presence and absence of chemotherapy was also enhanced when co-cultured with primary meningeal cells (Figure 1C). These results show that meningeal cells protect leukemia cells from the effects of cytotoxic chemotherapy.

**Figure 1.**
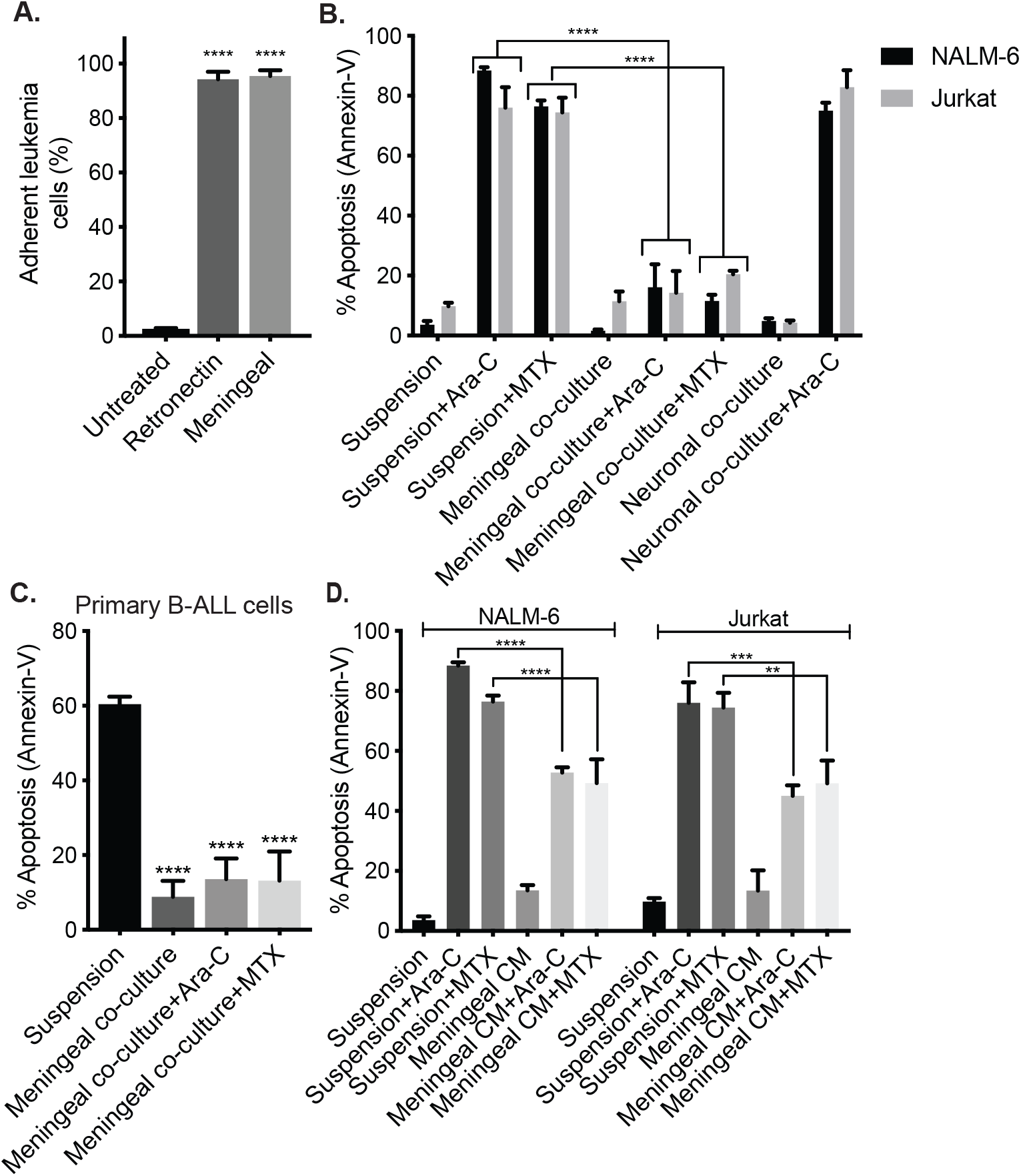
Leukemia cells exhibit increased chemoresistance when co-cultured with meningeal cells. (A) Percent of NALM-6 leukemia cells adherent to primary human meningeal cells, retronectin (recombinant fibronectin fragment) positive control, or non-tissue culture treated well after 2 hours. (B-C) NALM-6 and Jurkat leukemia cells (B) or primary B-cell ALL cells (C) cultured in suspension or adherent to CNS-derived cells (primary human meningeal cells or the HCN-2 neuronal cell line) were treated with cytarabine 500 nM or methotrexate 500 nM for 48 hours and then apoptosis measured using annexin-V and viability staining and flow cytometry. (D) NALM-6 and Jurkat cells were cultured in either regular media or meningeal conditioned media (CM) and treated with cytarabine 500 nM or methotrexate 500 nM for 48 hours before apoptosis was measured using annexin-V staining and flow cytometry. For all graphs, data are the mean +/- SEM from three independent experiments and *P*: **, <0.01, ***, <0.001, ****, <0.0001 by ANOVA.

We then used conditioned media from primary human meningeal cells to test whether direct cell-cell contact is required for meningeal-mediated leukemia chemoresistance. As shown in Figure 1D, meningeal conditioned media conferred moderate chemoresistance on leukemia cells, but to a lesser extent than co-culture. Together these data suggest that meningeal-mediated leukemia chemoresistance is primarily dependent upon cell-cell contact with smaller contributions from a soluble factor(s) secreted by meningeal cells.

### Meningeal cells shift the apoptotic balance toward survival in leukemia cells

In order to define the mechanism of meningeal-mediated leukemia chemoresistance we next examined the effect of co-culture on the apoptosis pathway in leukemia cells. In agreement with the annexin-V results (Figure 1B), co-culture of leukemia cells with primary meningeal cells attenuated apoptosis caused by cytarabine or methotrexate treatment as determined by both caspase 3/7 activity and measurement of leukemia cell mitochondrial potential with the dye TMRE (Figure 2A-B and Supplemental Figure 3). Next, we used an apoptosis antibody array to identify changes in the expression of multiple apoptosis family proteins in leukemia cells co-cultured with meningeal cells relative to suspension. In particular, the levels of several pro-apoptotic proteins (BID, caspase 3 & 8) decreased in leukemia cells co-cultured with meningeal cells (Figure 2C). However, assessing how the dynamic levels, activities, and complex interactions of these and other BCL-2 family of pro-and anti-apoptotic proteins (BH3 proteins) integrate to regulate the overall apoptotic balance is experimentally challenging. To try and capture the overall effect of the meninges on leukemia apoptotic balance, we then utilized BH3 profiling, a functional apoptosis assay that uses the response of mitochondria to perturbations by BH3 domain peptides, such as BIM, to predict the degree to which cells are primed to undergo apoptosis by the mitochondrial pathway^37,38^. Importantly, decreased mitochondrial priming following drug treatment has been shown to be highly predictive of chemotherapy resistance *in vitro* and *in vivo*^37,38^. Accordingly, we found that leukemia cells co-cultured with meningeal cells exhibited increased cytochrome C retention upon exposure to the BIM peptide compared to leukemia cells in suspension (Figure 2D). These data suggest that leukemia cells co-cultured with meningeal cells are significantly less primed to undergo apoptosis through the mitochondrial pathway than the same leukemia cells in suspension.

**Figure 2.**
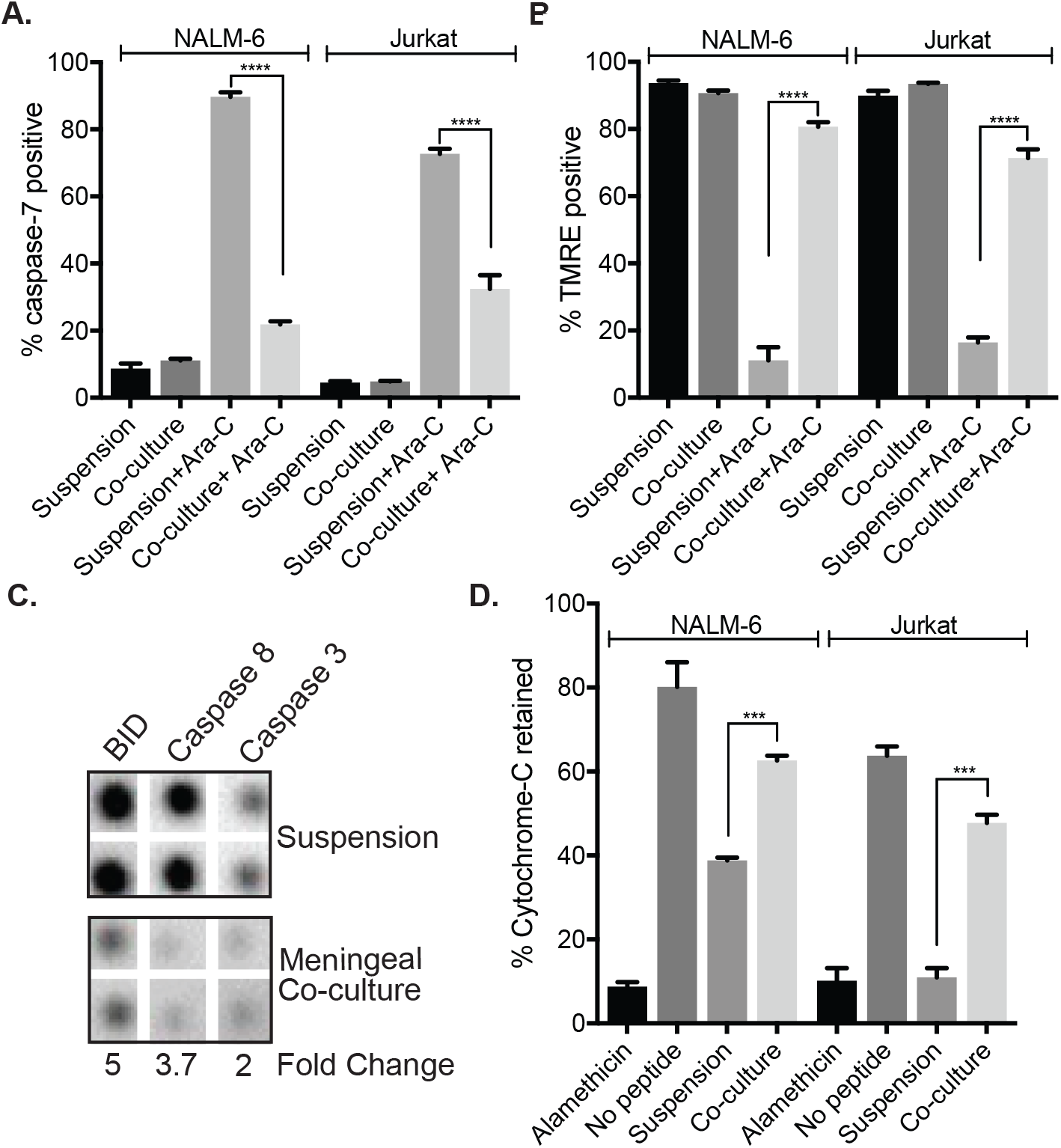
Meningeal cells tilt the apoptotic balance of leukemia cells toward survival. (A-B) NALM-6 and Jurkat leukemia cells cultured in suspension or adherent to meningeal cells were treated with cytarabine 500 nM for 48 hours and caspase-7 activity (A) and TMRE staining (B) assessed by flow cytometry. (C) NALM-6 leukemia cells were grown in suspension or adherent to meningeal cells for 48 hours. ALL cells were then isolated and lysed. Protein lysate was used to probe a Human Apoptosis Antibody Array (Abcam). Representative portions of the membrane are shown. Relative protein expression was calculated after normalization using the IgG positive control. (D) BH3 profiling was performed using the BIM peptide on NALM-6 and Jurkat leukemia cells grown in suspension or adherent to meningeal cells for 24 (NALM-6) or 48 hours (Jurkat). Cytochrome c retention was measured by flow cytometry. Alamethicin is a peptide antibiotic that permeabilizes the mitochondria membrane and serves as a positive control. For all graphs, data are the mean +/- SEM from three independent experiments and *P*: **, <0.01, ***, <0.001, ****, <0.0001 by ANOVA.

### Meningeal cells increase leukemia quiescence

We also examined the effect of the meninges on leukemia cell cycle and quiescence. As shown in Figure 3A-C, leukemia cells co-cultured with primary meningeal cells are less proliferative as shown by decreased Ki-67 staining, significantly decreased S phase, and increased G0/G1 phase. We next used Hoechst-Pyronin Y staining to better distinguish between G0 and G1 phases^29^. As shown in Figure 3D-F, leukemia cells in co-culture with primary meningeal cells exhibited increased G0 phase, indicative of quiescence, relative to leukemia cells in suspension. Glucose uptake also diminished in leukemia cells co-cultured with meningeal cells, further supporting that these leukemia cells are quiescent (Supplemental Figure 4).

**Figure 3.**
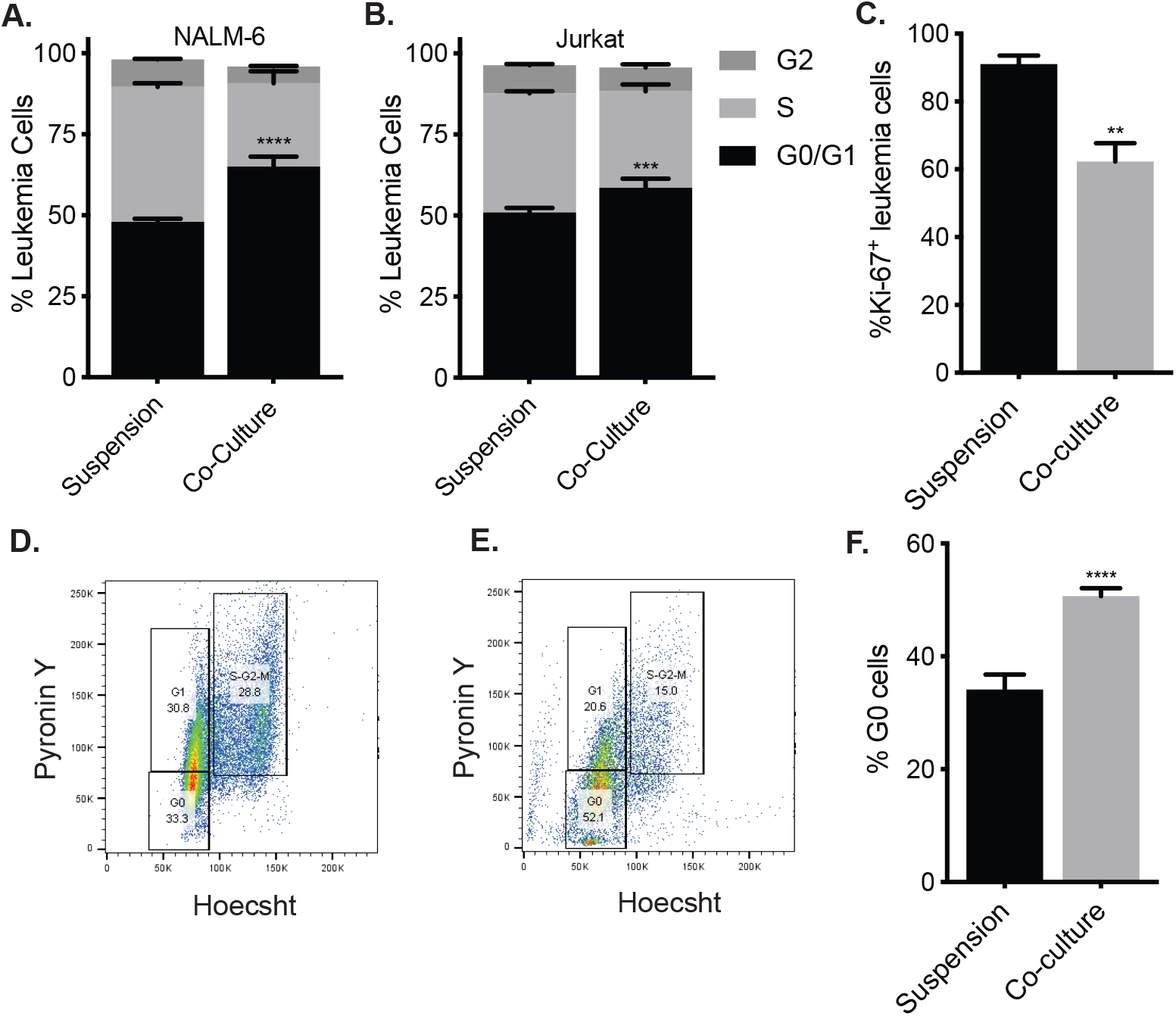
Meningeal cells increase leukemia quiescence *in vitro*. (A-B) NALM-6 and Jurkat leukemia cells cultured in suspension or adherent to primary meningeal cells for 48 hours were assessed for cell cycle and proliferation using a Click-iT Plus EdU kit and flow cytometry. (C-F) NALM-6 leukemia cells cultured in suspension or adherent to primary meningeal cells for 48 hours were stained for Ki-67 (C) or Hoechst-Pyronin Y (D-F) and analyzed by flow cytometry. Representative flow plots for Hoechst-Pyronin Y staining are shown (D-E). For all graphs, *P*: *, <0.05, **, <0.01, ***, <0.001, ****, <0.0001 by ANOVA.

We then examined cell cycle and quiescence in leukemia cells isolated from the meninges of xenotransplanted mice. As shown in Figure 4A-C, leukemia cells isolated from the meninges of transplanted mice exhibited increased G0/G1 phase and decreased Ki-67 staining relative to leukemia cells isolated from peripheral blood or bone marrow. In order to assess whether the meninges harbor long-term quiescent leukemia cells, we labeled leukemia cells with a fluorescent membrane dye (DiR or DiD) that is retained in dormant, non-cycling cells but is diluted to undetectable levels within 3-5 generations in proliferating leukemia cells^28,30–33^. Dye-labeled leukemia cells were then transplanted into immunodeficient mice. After systemic leukemia development in ~4 weeks, the mice were euthanized and meninges harvested. Within the meninges we detected low levels of dye-retaining, quiescent leukemia cells by flow cytometry (<1% of total leukemia population; Figure 4D-E). Further supporting that these dye-retaining leukemia cells are quiescent, the dye-retaining leukemia cells from the meninges exhibited low expression of the proliferation marker Ki-67 (Figure 4F). We also detected these quiescent leukemia cells within the meninges using confocal microscopy (Figure 4G-I).

**Figure 4.**
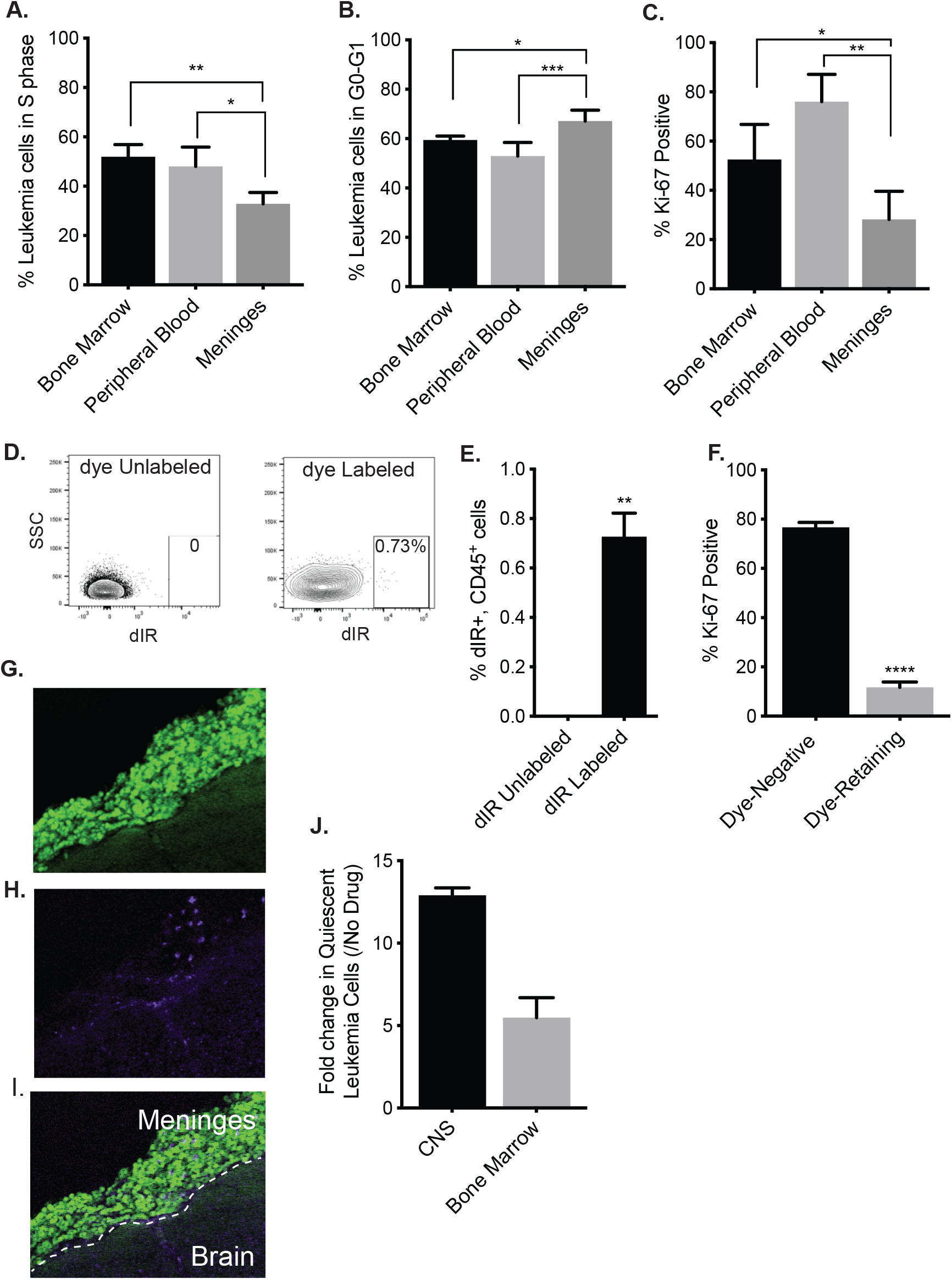
The meninges harbor quiescent and chemoresistant leukemia cells *in vivo*. (A-C) Mice were transplanted with NALM-6 leukemia cells (2×10^6^ cells; N=5 per group). After 4 weeks, mice were injected IV with Click-iT EdU cell cycle reagent prior to harvesting tissues (meninges, bone marrow, and peripheral blood) and assessing for cell cycle (A-B) and the proliferation marker Ki-67 (C) by flow cytometry. (D-F) Mice were transplanted with membrane dye-labeled (DiD) or control, unlabeled NALM-6 leukemia cells (2×10^6^ cells) and then euthanized 4 weeks later (N=5). Ki-67 negative, quiescent (dye-positive) ALL cells were identified and quantitated (D-F) in the meninges by flow cytometry. Representative flow cytometry plots are shown in (D). (G-I) Confocal microscopy images showing total (G; green), dye-retaining, quiescent (H; purple), and overlay (I) of leukemia cells detected in the meninges of mice 4 weeks after transplantation with dye-labeled (DiR) NALM-6 cells also expressing GFP. (J) Mice were transplanted with dye-labeled human NALM-6 ALL cells (2×10^6^; N=5 per group). After 3 weeks, the dye-positive and dye-negative ALL cells were measured and quantitated in the meninges and bone marrow by flow cytometry before and after treatment with cytarabine 50 mg/kg x 3 days. A relative increase in the dye-positive leukemia cells after cytarabine treatment is consistent with dye-positive leukemia cells being chemoresistant relative to the dye-negative leukemia cells. For all graphs, *P*: *, <0.05, **, <0.01, ***, <0.001, ****, <0.0001 by ANOVA or t-test.

We next treated these xenotransplanted mice with cytarabine and measured the percentage of dye-retaining leukemia cells in the meninges by flow cytometry. Cytarabine was given at a dose previously shown to result in plasma levels in the range of human high-dose cytarabine regimens that cross the blood-brain barrier^27,39^. Cytarabine, relative to PBS, also significantly reduced the CNS leukemia burden in xenotransplanted mice and provides functional data that cytarabine is crossing the blood-brain barrier in our murine experiments. (Supplemental Figure 5). Moreover, the relative increase in dye-retaining leukemia cells after cytarabine treatment is consistent with these cells having increased chemoresistance compared to the dye-negative, proliferating leukemia cells (Figure 4J). Together these data suggest the meninges harbor quiescent leukemia cells that exhibit chemoresistance.

### Meningeal-mediated leukemia chemoresistance is a reversible phenotype

We then tested whether removal of leukemia cells from co-culture with meningeal cells restored chemosensitivity. Leukemia cells were dissociated from meningeal cells, purified with CD19 (NALM-6) or CD3 (Jurkat) magnetic beads, and placed back in suspension. As shown in Figure 5A, leukemia cells removed from co-culture exhibited similar sensitivity to methotrexate and cytarabine as leukemia cells in suspension. Similarly, leukemia cells grown in suspension after isolation from the meninges of xenotransplanted mice also exhibited sensitivity to cytarabine (Figure 5B). Further supporting these results, leukemia cells placed back into suspension after co-culture reverted back to baseline cell cycle and apoptosis characteristics (Figure 5C-E). These results suggest that drugs that disrupt adhesion between leukemia and meningeal cells may restore leukemia chemosensitivity in the CNS niche.

**Figure 5.**
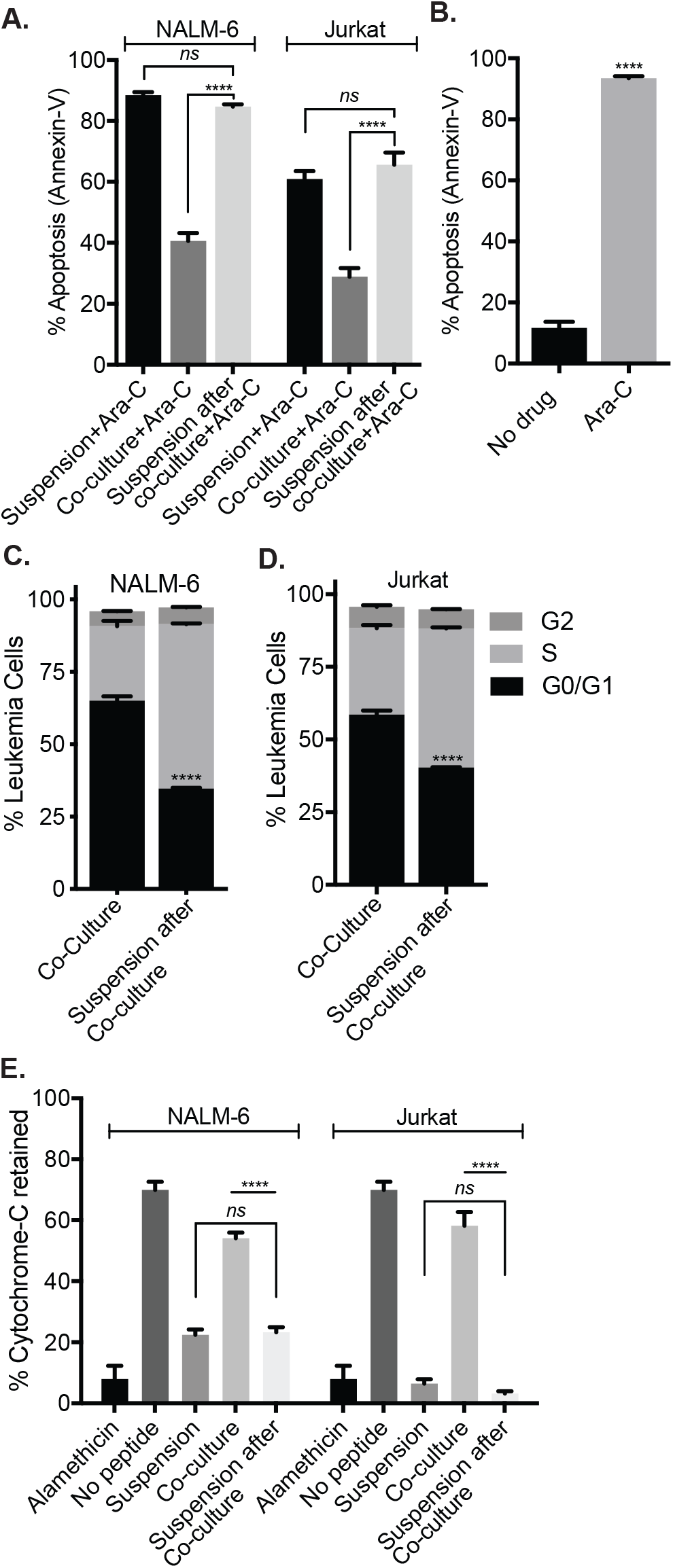
The effects of the meninges on leukemia cells are reversible. (A) NALM-6 and Jurkat cells grown in suspension, co-cultured with meningeal cells, or isolated from co-culture and placed back in suspension were treated with Ara-C 500 nM and apoptosis assessed by annexin-V staining and flow cytometry. (B) NALM-6 leukemia cells were isolated from the meninges of xenotransplanted mice and then grown in suspension for 48 hours prior to treatment with cytarabine 500 nM and assessment of viability using annexin-V staining and flow cytometry. (C-D) NALM-6 (C) and Jurkat (D) leukemia cells were co-cultured with meningeal cells or isolated from co-culture and placed back in suspension and then cell cycle assessed using Click-iT Plus EdU cell cycle reagent and flow cytometry. (E) BH3 profiling was performed using the BIM peptide on NALM-6 and Jurkat leukemia cells co-cultured with meningeal cells or placed back into suspension after co-culture with meningeal cells. Cytochrome c retention was measured by flow cytometry. For all graphs, data are the mean +/- SEM from three independent experiments and *P*: ****, <0.0001 by ANOVA. *ns*, not significant.

### Overcoming meningeal-mediated leukemia chemoresistance by disrupting adhesion

Given that meningeal-mediated leukemia chemoresistance is a reversible phenotype, we next identified several cell adhesion inhibitors that disrupted leukemia-meningeal adhesion in co-culture (Supplemental Figure 6A). We selected Me6TREN for further testing based upon its effectiveness in co-culture, small molecular weight (increasing likelihood for CNS penetration), tolerability in mice, lack of prior testing in leukemia, and likely multifactorial mechanism of action^25,26^. As shown in Figure 6A, Me6TREN significantly disrupted the adhesion of both NALM-6 and Jurkat leukemia cells to primary meningeal cells. This effect of Me6TREN on adhesion was not due to toxicity to leukemia or meningeal cells (Supplemental Figure 6B). We then quantitated non-adherent leukemia cells in the cerebral spinal fluid (CSF) of xenotransplanted mice after treatment with either Me6TREN or PBS control in order to measure disruption of leukemia adhesion *in vivo*. As shown in Figure 6B, Me6TREN treated mice showed a modest, but significant, increase in leukemia cells in the CSF, consistent with the co-culture experiment. Moreover, by disrupting leukemia adhesion, Me6TREN significantly attenuated leukemia chemoresistance in co-culture with meningeal cells (Figure 6C). We next assessed the *in vivo* ability of Me6TREN to enhance the efficacy of cytarabine in treating leukemia in the meninges. We tested NALM-6, Jurkat, and primary B-ALL leukemia cells with dosing regimens shown in Supplemental Figure 7. In all cases, Me6TREN significantly enhanced the efficacy of cytarabine in reducing the number of viable leukemia cells in the meninges (Figure 6D-G). Finally, despite Me6TREN disrupting the bone marrow hematopoietic niche^25^, mice receiving cytarabine/Me6TREN or cytarabine alone exhibited comparable hematologic toxicities (Supplemental Figure 8).

**Figure 6.**
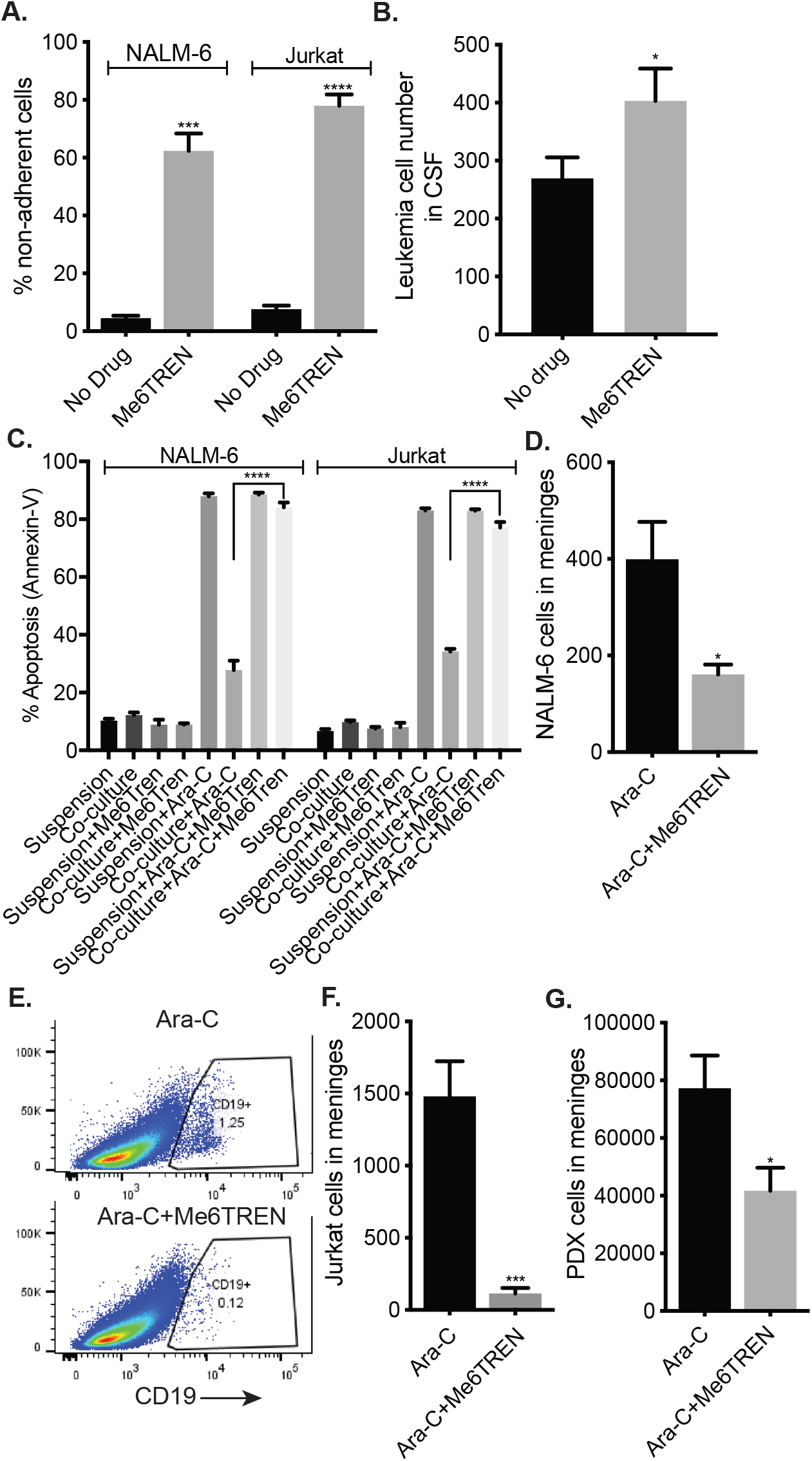
Me6TREN disrupts the meningeal-leukemia niche and attenuates leukemia chemoresistance *in vitro* and *in vivo*. (A) NALM-6 leukemia cells were co-cultured with meningeal cells in the presence or absence of Me6TREN 100 μM. After 24 hours, non-adherent leukemia cells were removed and quantitated. (B) Mice were transplanted with NALM-6 leukemia cells (2×10^6^ cells; N=5 per group). After 16 days, mice were treated with Me6TREN 10 mg/kg subcutaneous or PBS control x 3 days. After an additional 24 hours, 5-10 μL of cerebral spinal fluid was removed from the cisterna magna and leukemia cells quantitated using flow cytometry and count beads. (C) Leukemia cells in suspension or co-cultured with primary meningeal cells were treated with Me6TREN +/- cytarabine 500 nM and after 48 hours viability assessed with annexin-V staining and flow cytometry. (D-G). Mice transplanted with NALM-6 (2×10^6^ cells; N=5 per group; D & E), Jurkat (2×10^6^ cells; N=5 per group; F), or primary B-ALL (PRoXe Sample CBAB-62871-V1; 1×10^6^ cells; N=5 per group; G) leukemia cells were treated with cytarabine (50 mg/kg intraperitoneal) or cytarabine + Me6TREN (10 mg/kg subcutaneous). 48 hours after completing therapy mice were euthanized, cardiac perfused, meninges isolated and dissociated, stained with human CD19 (NALM-6 & PDX) or CD3 (Jurkat)antibody, and leukemia cells quantitated by flow cytometry and counting beads (D, F, G). Representative flow plots for NALM-6 are shown in (E). For all graphs, *P*: *, <0.05, **, <0.01, ***, <0.001, ****, <0.0001 by ANOVA or t-test.

## Discussion

Protection from leukemia relapse in the CNS is crucial to long-term survival and quality of life for leukemia patients^1–3^. One strategy for developing novel CNS-directed therapies has focused on identifying, and potentially targeting, the factors that facilitate leukemia cell trafficking to the CNS from the bone marrow^40–49^. At the same time, additional evidence suggests that the ability of leukemia cells to infiltrate the CNS is a general property of most leukemia cells and not restricted to rare clones that acquire a metastatic phenotype^14,15^. Accordingly, we sought to address the question of how leukemia cells adapt to this unique niche and escape the effects of chemotherapy after infiltrating the CNS. We found that within the CNS leukemia cells primarily localize to the meninges and that parenchymal involvement by leukemia was a rare finding. This observation is in agreement with a larger body of literature demonstrating that leukemia xenografts accurately model the anatomic distribution of leukemia observed within the CNS of humans^11–14^. As a result, we focused our work on the meninges. However, it is certainly possible, and perhaps even likely, that other cells or tissues within the CNS, such as the choroid plexus, may also impact leukemia biology^17,20^. This may be analogous to the bone marrow microenvironment in which distinct niches (endosteal, vascular) exert unique effects on hematopoietic stem and leukemia cells^50^.

We then used co-culture and *in vivo* xenotransplantation approaches to further characterize the effects of the meninges on leukemia biology. We found that the meninges enhance leukemia resistance to cytarabine and methotrexate, the primary drugs currently used in the treatment of CNS leukemia, by altering the apoptotic balance in leukemia cells to favor survival and increasing leukemia quiescence ^1,2^. Quiescence allows cancer cells to escape cytotoxic chemotherapy and has been shown to be critical for leukemia relapse and stem cell biology^28,51,52^. In agreement, it was previously shown that high Mer kinase expressing, t(1;19) leukemia cells co-cultured with CNS-derived cells exhibit G0/G1 cell cycle arrest, suggestive of dormancy or quiescence, as well as methotrexate resistance^16^.

To define the mechanism by which the meninges exert these effects on leukemia biology, we also tested the ability of meningeal conditioned media to enhance leukemia chemoresistance. While meningeal conditioned media partially attenuated the sensitivity of leukemia cells to chemotherapy, the effect was significantly less than when leukemia cells were in direct contact with meningeal cells. This result supports a model in which leukemia chemoresistance is primarily dependent upon direct interactions between the leukemia and meningeal cells with smaller contributions from a soluble factor(s) secreted by the meningeal cells. Together these results further support that the pathophysiology of CNS leukemia and relapse is more complex than simply the ability of leukemia cells or chemotherapy to access the restricted CNS microenvironment and complement other extensive lab and clinical data demonstrating that cell-autonomous factors play an essential role in leukemia biology^50,53,54^.

Importantly, we also found that meningeal-mediated leukemia chemoresistance was a reversible phenotype. Leukemia cells removed from co-culture with meningeal cells or the meninges of xenotransplanted mice reverted back to baseline cell cycle, quiescence, apoptosis balance, and sensitivity to methotrexate and cytarabine. Ebinger *et al*. recently identified a similar population of relapse-inducing ALL cells within the bone marrow that exhibited dormancy, stemness, and treatment resistance^30^. However, similar to our results, these therapeutically adverse properties were reversed when these leukemia cells were dissociated from the bone marrow microenvironment.

We then identified drugs capable of disrupting leukemia-meningeal adhesion. In addition to identifying several drugs that inhibit canonical cell adhesion targets, we also found that Me6TREN, a novel hematopoietic stem cell (HSC) mobilizing compound, also disrupted leukemia-meningeal adhesion *in vitro* and *in vivo*. Moreover, Me6TREN enhanced the efficacy of cytarabine in treating CNS leukemia in xenotransplanted mice. *In vivo* efficacy against two leukemia cell lines with distinct immunophenotypes (T and B-cell) and a primary B-ALL PDX supports the possibility that drugs, or biologic agents, that target leukemia–niche interactions may exhibit broader specificity than mutation-specific therapeutics that are limited by the genetic heterogeneity of leukemia. The mechanism by which Me6TREN disrupts the leukemia niche is not known. However, Me6TREN mobilizes mouse hematopoietic stem/progenitor cells from the bone marrow through effects on several cell adhesion and migration pathways including MMP-9, SDF-1/CXCR4, and VECAM/VLA-4^25^. As these adhesion pathways also contribute to the bone marrow leukemia niche, it is possible that Me6TREN may exhibit efficacy in other leukemia niches ^53,55^. However, if upon further testing Me6TREN only disrupts leukemia-meningeal adhesion, Me6TREN could still be effective in treating isolated CNS relapses or for use in upfront therapy to reduce the risk for CNS relapse.

Given the complexity of cell-cell interactions, this multi-faceted mechanism of action of Me6TREN may provide an advantage relative to other niche-disrupting agents being developed for leukemia therapy that target a single mechanism of retention or adhesion (i.e. inhibitors of SDF-1 or a single extracellular adhesion factor)^56^. An alternative approach to overcome the complexity and redundancy of cellular adhesion would be to combine multiple inhibitors that target different adhesion molecules (Supplemental Figure 6A).

A potential risk of combining Me6TREN, or other niche-disrupting agents, and chemotherapy is that normal HSCs mobilized into the circulation by ME6TREN may be sensitized to chemotherapy with resulting marrow aplasia or delayed hematopoietic recovery. However, we found that mice receiving cytarabine/Me6TREN or cytarabine alone exhibited comparable hematologic toxicities. Moreover, preclinical and clinical studies combining other HSC mobilizing agents, such as AMD3100 or G-CSF, with chemotherapy also have shown an acceptable toxicity profile^57–59^. Another potential limitation to this approach of disrupting leukemia cell adhesion to the niche is that soluble factors secreted by the niche can also interact with non-adherent leukemia cells and impact leukemia biology, as we saw with meningeal conditioned media ^46,60^. Combination therapies that target both adhesion and secreted factors may be the most efficacious in treatment of CNS leukemia.

In summary, this work demonstrates that the meninges enhance leukemia chemoresistance in the CNS, elucidates mechanisms of CNS relapse beyond the role of the blood-brain barrier, and identifies niche disruption as a novel therapeutic approach for enhancing the ability of chemotherapy to eradicate CNS leukemia.

## Acknowledgements

This work was supported in part by the Children’s Cancer Research Fund (PMG), the Timothy O’Connell Foundation (PMG), and an American Cancer Society Institutional Research Grant (PMG). PB was partially supported by NIH Training Grant T32 CA099936. We thank Dr. Michael Farrar for providing mouse BCR/ABL p190 leukemia cells. We also thank Dr. Mark Sanders (University of Minnesota Imaging Center) for providing expert assistance with confocal microscopy and sample preparation. This work utilized the University of Minnesota Masonic Cancer Center shared flow cytometry and comparative pathology resources and the Hematological Malignancy Tissue Bank, which are supported in part by NCI 5P30CA077598-18, Minnesota Masonic Charities, and the Killebrew-Thompson Memorial Fund.

## Authorship

Contributions: L.M.J. and M.E. designed experiments, analyzed data, and prepared figures. P.B., J.M., and K.J. performed experiments and analyzed data. P.M.G. designed the study, oversaw the laboratory investigations, and wrote the manuscript. All authors reviewed, edited and approved the final version of the manuscript.

## Competing Interests Disclosure

The authors declare no competing financial interests.

