## Supplemental Materials for "Disrupting the leukemia niche in the central nervous system attenuates leukemia chemoresistance"

***Supplemental Methods***

**Immunohistochemistry:** NSG and C57BL/6 mice were transplanted as described with human and mouse leukemia cells, respectively. At signs of systemic leukemia, mice were euthanized and cardiac perfused with PBS followed by paraformaldehyde 4%. The brains and meninges were removed, embedded in paraffin, processed, and stained with either human CD19 or mouse CD45.1 antibody.

**Glucose Uptake:** The effect of co-culture on leukemia glucose uptake was measured using the Glucose Uptake Assay Kit (Abcam) according to the manufacturer's instructions. In brief, leukemia cells were cultured in regular media in suspension or adherent to meningeal cells. After 48 hours, the media was switched to RPMI supplemented with FBS 0.5% and a fluorescent glucose analog. After one hour, leukemia adhesion was disrupted by trypsinization and manual pipetting. The leukemia cells were then washed, suspended in Analysis Buffer, and analyzed by flow cytometry to determine median fluorescent intensity (MFI).

**Mouse Hematopoiesis:** Baseline complete blood counts were obtained on C57 BL/6 mice using a Hemavet Blood Analyzer (Drew Scientific). Mice were then treated with cytarabine (50 mg/kg x 5 day), cytarabine (50 mg/kg x 5 day)/Me6TREN (10 mg/kg x 5 days), or untreated. Complete blood counts were then measured at days 5, 10, and 20.

**Conditioned Media:** Primary meningeal cells were cultured until ~60-70% confluent and then fresh media was added. After an additional 48-72 hours, media was collected, centrifuged at 2000 rpm for 10 minutes to remove any cellular debris, and then filtered through as 0.45 μm filter (Millipore).

**BH3 Profiling:** BH3 profiling was performed as described(34,35). Leukemia cells labeled with CellTrace dye were grown in suspension or co-culture with primary meningeal cells for 24-48 hours and then suspended in membrane extraction buffer containing digitonin 100 μg/mL (Sigma-Aldrich). The leukemia cells were then incubated with the BIM peptide 50 μM (GenScript) for 90 minutes. After incubation, leukemia cells were fixed with paraformaldehyde 4%, permeabilized, stained for cytochrome c (eBioscience), and analyzed by flow cytometry.

**Adhesion assay:** 1-1.5 x $10^5$ leukemia cells were added to ~80% confluent meningeal cells. Simultaneously, Me6Tren 100 $\mu$M was added to co-culture media. After 24-48 hours, the media, including non-adherent leukemia cells, was gently removed and viable leukemia cells quantitated by both manual counting with a hemocytometer and flow cytometry with Count Bright Absolute counting beads (ThermoFisher).

*Supplemental Figure Legends*

**Supplemental Figure 1. Leukemia cells localize to the meninges within the CNS.** (A) Schematic illustrating the xenotransplantation model. (B-C) NSG or C57BL/6 mice were transplanted with human NALM-6 ($2x10^6$ cells; N=5 mice) or murine BCR/ABL p190 (3000 cells; N=5 mice) leukemia cells, respectively. After 3 weeks (NALM-6 mice) or 10 days (BCR/ABL p190 mice), the mice were euthanized and cardiac perfused with PBS followed by fixative. IHC for human CD19 (B) or mouse CD45.1 (C) was then used to identify leukemia cells within the CNS. Leukemia cells stain brown. Original magnification x20 for panels B-C.

**Supplemental Figure 2. Chemotherapy does not significantly affect the viability of primary human meningeal cells.** Primary human meningeal cells at ~80% confluence were treated with either cytarabine (Ara-C) 500 nM, methotrexate (MTX) 500 nM, or mock treated for 48 hours and then viability assessed with annexin-V staining and flow cytometry. *ns*, not significant.

**Supplemental Figure 3: Meningeal cells tilt the apoptotic balance of leukemia cells toward survival.** (A-B) NALM-6 and Jurkat leukemia cells cultured in suspension or adherent to meningeal cells were treated with methotrexate 500 nM for 48 hours and caspase-7 activity (A) and TMRE staining (B) assessed by flow cytometry. For both graphs, data are the mean +/- SEM from three independent experiments and *P*: ****, <0.0001 by ANOVA.

**Supplemental Figure 4. Co-culture with meningeal cells decreases glucose uptake by leukemia cells.** NALM-6 and Jurkat leukemia cells were cultured in suspension or adherent to meningeal cells for 48 hours prior to treatment with a fluorescent glucose analog. Uptake of the analog was then determined by measuring the median fluorescent intensity (MFI) by flow cytometry. *P*: **, <0.01, ***, <0.001 by t-test.

**Supplemental Figure 5: Systemic cytarabine reduces the leukemia burden in the CNS.** Mice transplanted with NALM-6 leukemia cells ($2x10^6$ cells; N=5 per group) were treated with cytarabine (50 mg/kg intraperitoneal) x 5 days or PBS. 48 hours after completing therapy mice were euthanized, cardiac perfused, meninges isolated and dissociated, stained with human CD19 antibody, and leukemia cells quantitated by flow cytometry. *P*: ***, <0.001 by t-test.

**Supplemental Figure 6: Identification of compounds that disrupt the adhesion of leukemia and meningeal cells.** (A) NALM-6 leukemia cells were added to primary meningeal cells in the presence of A205804 1 μM (E-selectin and ICAM-1 inhibitor,) AMD3100 100 μM (CXCR4 antagonist), KF38789 200 μM (P-selectin inhibitor) or RGD 20 μM (integrin inhibitor). After 24 hours, the media, and any non-adherent leukemia cells were removed, and leukemia cells quantified. *P*: ****, <0.0001 by ANOVA. (B) NALM-6 leukemia cells and primary meningeal cells were treated with Me6TREN 100 μM or mock treated for 48 hours and then viability assessed with annexin-V staining and flow cytometry. Viability relative to the mock treated cells is plotted.

**Supplemental Figure 7: Different drug treatment regimens tested in vivo*.*** Schematics illustrating the different *in vivo* dosing regimens used for NALM-6, Jurkat, and primary B-ALL PDX leukemia cells. Treatment start dates and duration of cytarabine treatment varied between the xenografts because of differences in the rate of CNS engraftment and baseline cytarabine sensitivity. In all experiments, the dose of cytarabine was 50 mg/kg IP daily and Me6TREN 10 mg/kg sc daily.

**Supplemental Figure 8: Effect of Me6TREN and chemotherapy on hematopoiesis.** C57 BL/6 mice were untreated (black), treated with cytarabine (50 mg/kg x 5 days; red) or cytarabine/Me6TREN (10 mg/kg x 5 days; blue) and then blood counts were measured on specified days. White blood cell (A) and platelet (B) counts are shown for individual mice.

***Supplemental Table Legend***

**Supplemental Table 1*.*** Multiple human leukemia cell lines cultured either in suspension or adherent to primary human meningeal cells were treated with cytarabine or methotrexate for 48 hours and then apoptosis was measured using annexin-V staining and flow cytometry. In the table, + signifies that $P < 0.05$ when comparing leukemia chemosensitivity in suspension versus in co-culture.

### Supplemental Figure 1

A.

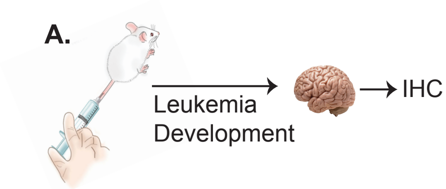


B.

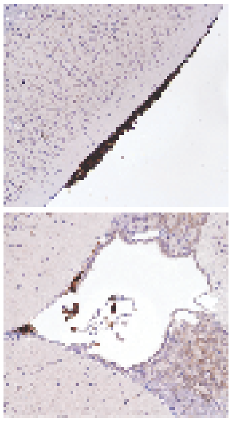

C.

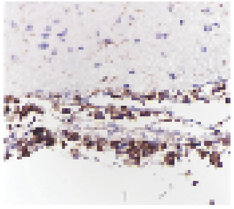

**Supplemental Figure 2**

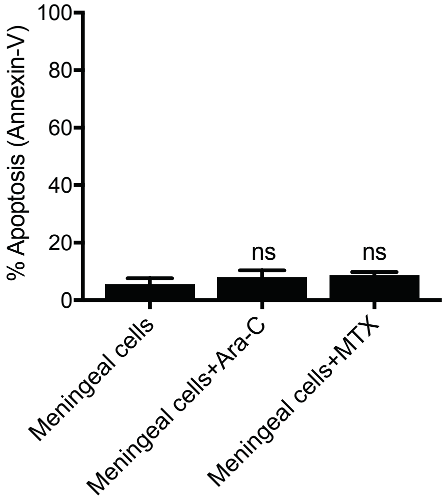

### Supplemental Figure 3

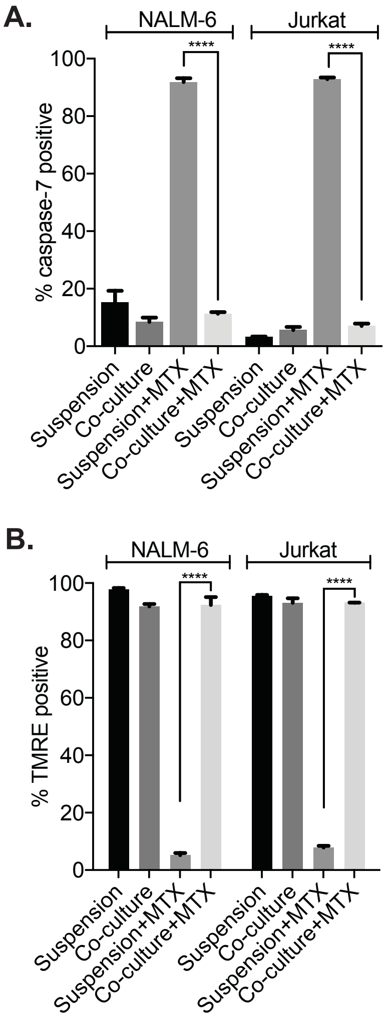

**Supplemental Figure 4**

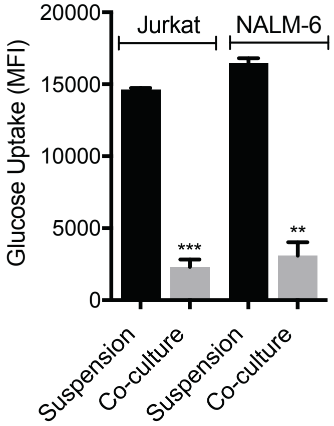

**Supplemental Figure 5**

**A.**

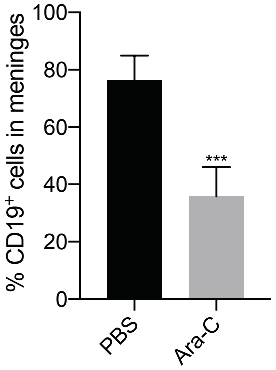

### Supplemental Figure 6

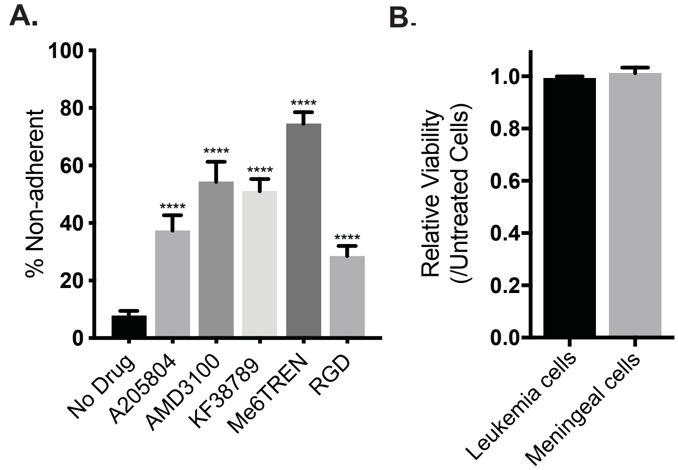

**Supplemental Figure 6**

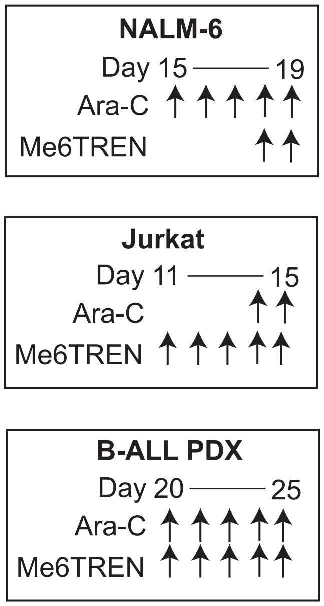

**Supplemental Figure 7**

**A.**

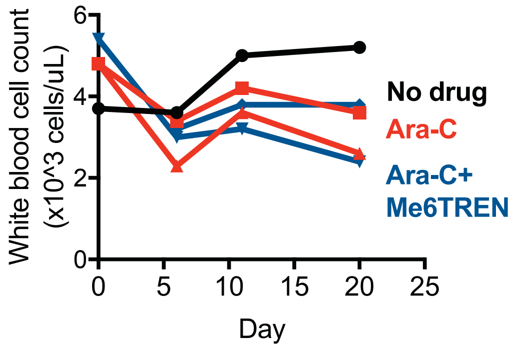


**B.**

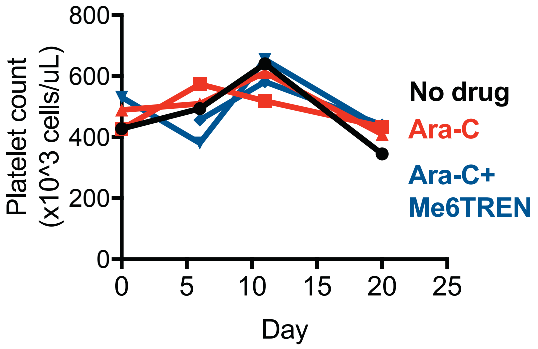
